# A CBF–SA module links wound-induced evaporative cooling to tissue repair in plants

**DOI:** 10.1101/2025.05.23.655667

**Authors:** Joseph Balem, Chenjiao Tan, Nathália Cássia Ferreira Dias, Mariah Arnold, Sorrel Tran, Paul M. Severns, Paulo J.P.L. Teixeira, Changying Li, Li Yang

## Abstract

Repairing damaged tissues is essential for the survival of all organisms. In plants, tissue injury rapidly triggers defense and repair programs. However, the molecular mechanisms linking early injury cue to the later stages of wound repair remain unclear. Here, we show that wounding of Arabidopsis leaves induces localized low temperature at the injury site, likely caused by evaporative cooling, which is accompanied by an activation of cold-responsive genes. Using thermal imaging combined with computer vision and deep learning, we developed a workflow to monitor the dynamics of wound healing in a quantitative, non-invasive and real-time manner. Mechanistically, we show that C-repeat Binding Factor (CBF) transcription factors are required for the activation of injury-associated cold response and downstream salicylic acid (SA) signaling. The CBF–SA module promotes lignin deposition and wound repair. Together, these findings reveal a link between a wound-induced biophysical cue and the tissue repair program.

## Introduction

Responding to and repairing tissue damage is essential for an organism’s survival. In nature, wounds to aboveground plant tissues can be caused by abiotic sources such as hail and strong winds, as well as biotic interactions such as herbivore feeding. Natural developmental processes, such as lateral root emergence and leaf shedding, also leave injured tissues. In addition, wounds are purposefully created in agricultural and horticultural practices such as grafting, pruning, harvesting and tissue regeneration. Before wounds are fully healed, their presence may not only impair proper organ functioning, but can become entry points for microbial pathogens, resulting in plant disease. Therefore, wound healing is an important adaptive response to tissue damage that occurs throughout a plant’s lifespan.

The wound response in *Arabidopsis thaliana* involves a rapid and dynamic network of signaling pathways that limits damage, suppresses pathogen invasion, and promotes tissue repair (Schilmiller & Howe, 2005; Ikeuchi *et al*., 2020; Vega-Muñoz *et al*., 2020). When injured, damaged cells immediately release damage-associated molecular patterns (DAMPs), such as cell wall fragments and cytosolic content (e.g. ATP, Glutathione) (Tanaka & Heil, 2021). The perception of DAMPs by neighboring cells triggers a cascade of intracellular signaling events, including calcium ion influx and the production of reactive oxygen species (ROS) (De Lorenzo *et al*., 2018; Tanaka & Heil, 2021). Mitogen-activated protein kinase (MAPK) cascades, such as MPK3 and MPK6, transmit damage signals to downstream responses (Beckers *et al*., 2009; Zhang & Zhang, 2022). Phytohormones, particularly jasmonic acid (JA), play key roles in modulating wound responses (Koo & Howe, 2009). The role of mechanical barriers, such as cuticle repair and suberization, contributes to sealing the wound site and preventing water loss or pathogen entry (Serrano *et al*., 2014; Rains *et al*., 2022). Thus, wound-triggered deposition of callose and lignin are usually considered as hallmarks of the healing process (Jacobs *et al*., 2003; Xu *et al*., 2024), although the threshold levels of callose and lignin required for effective wound sealing remain unclear.

Sensing and responding to temperature fluctuations is a crucial ability for plants adapting to a changing environment. C-repeat Binding Factor (CBF) transcription factors act as central regulator of cold response in Arabidopsis (Cook *et al*., 2004; Jia *et al*., 2016; Hwarari *et al*., 2022). Mutations in CBF genes lead to various defects related to sensing cold temperature including reduced freezing tolerance, altered flowering time, and reduced accumulation of osmoprotectants (Gilmour *et al*., 1998; Jia *et al*., 2016; Hwarari *et al*., 2022), which are acclimatization and adaptive responses to low temperature. CBFs bind to a C-repeat/Dehydration Responsive Element (CRT/DRE) motif in the promoters of genes responsive to cold stresses including a variety of proteins annotated as Cold Regulated genes (COR genes) (Stockinger *et al*., 1997; Liu *et al*., 1998; Park *et al*., 2015; Hwarari *et al*., 2022). In different temperature ranges, *COR* genes have multiple functions in stabilizing membrane phospholipids, maintaining hydrophobic interactions, ion homeostasis, and scavenging ROS (Artus *et al*., 1996; Steponkus *et al*., 1998; Puhakainen *et al*., 2004; Bozovic *et al*., 2013; Wang *et al*., 2017; Ortega *et al*., 2024).

While significant progress has been made in understanding the wound response in plants, a critical knowledge gap remains in elucidating the mechanisms that drive the later stages, particularly wound healing. Currently, there is a lack of robust, quantitative methods to measure and track wound healing dynamics, limiting our ability to connect early wound signaling events to the final healing step. Studies have demonstrated the potential of Deep Learning (DL) in diverse applications, including plant phenotyping (Murphy *et al*., 2024), stress detection (Islam *et al*., 2024b), and yield prediction (Jhajharia *et al*., 2023). Advances in DL, particularly convolutional neural networks (CNNs), have enabled precise and efficient segmentation and detection of leaves from complex images. For instance, U-Net and its variants have been widely used for leaf segmentation, achieving high accuracy in splitting leaves from complex backgrounds (Bhagat *et al*., 2022; Deng *et al*., 2023; Fu *et al*., 2023; Yi *et al*., 2023; Zhang *et al*., 2026 Zhang, 2026 #6200). Additionally, instance segmentation models such as Mask R-CNN have proven effective in parsing regions within individual leaves (Chowdhury *et al*., 2024; Huang *et al*., 2024; Islam *et al*., 2024a; Alfred *et al*., 2025), allowing for accurate quantification of leaf traits. These methods have significantly reduced the manual effort to extract features and improved the accuracy of image-based monitoring, making high-throughput phenotyping feasible. For instance, mask R-CNN was employed to segment plants in the thermal images, achieving an error ≤ 0.5 ℃ (Jiang *et al*., 2018). Similarly, Mask R-CNN was utilized to segment apples in RGB images. The segmentation masks were then mapped onto the thermal images to measure apple temperatures, achieving an error < 0.5 ℃ (Amogi *et al*., 2024).

Here we report that a DL-based workflow was capable of accurately monitoring wound-induced temperature differences. We present results demonstrating that wound-induced evaporative cooling is necessary and sufficient to induce a localized cold response. The key players in cold response, CBF transcription factors, were required for the activation of wound-induced cold response locally. In addition, CBFs also contribute to the wound-induced SA response. Mutations in both CBFs and SA biosynthesis/signaling genes led to delayed healing and reduced lignin deposition at wound sites. Based on these observations, we propose that CBF-mediated temperature sensing pathway may relay the wound-associated evaporative cooling signal to SA-governed wound repair in Arabidopsis.

## Results

### Cold-response genes were activated following wounding

To investigate the early transcriptional response to wounding, we analyzed the transcriptomic changes at 30 minutes and 60 minutes after cutting in detached *Arabidopsis* leaves. Compared to the control condition (time 0), a total of 6132 genes were differentially expressed at one or both time points. Specifically, 4522 genes were differentially expressed at 30 minutes, 5107 at 60 minutes, and 3497 genes were shared between the two time points. Hierarchical clustering grouped these wound-responsive genes into eight distinct clusters based on their expression profiles (Figure 1A, Supplementary Table 1). Clusters 1-6 comprised 3154 genes that were up-regulated in response to wounding, while Clusters 7 and -8 included 2978 genes that were down-regulated (Figure 1B). As expected, JA-responsive genes were prominently activated upon wounding (Supplementary Figure 1A). Interestingly, genes associated with cold response were significantly enriched among the up-regulated genes, particularly in Clusters 1, 2 and 6 (Supplementary Figure 1). Notably, Cluster 2 and 6 included well-known marker genes of the cold response (i.e., *COR6.6*, *RD29A*, *KIN1*, *COR47A*, *COR15A*, and *COR15B*) (Figure 1A and 1C), indicating that wounding activated a cold-response-like program in plants. To validate these findings, we cross-referenced our results with published transcriptomes from wounded Arabidopsis hypocotyls (Ikeuchi *et al*., 2017), leaf blade (Fiorucci *et al*., 2022) and petiole (Kilian *et al*., 2007; Liu *et al*., 2022). Consistent with our observations, cold-responsive marker genes were also up-regulated in these datasets (Supplementary Figure 1B). Together, these results suggest that activation of the cold-response pathway is associated with the Arabidopsis wound response.

**Figure 1:**
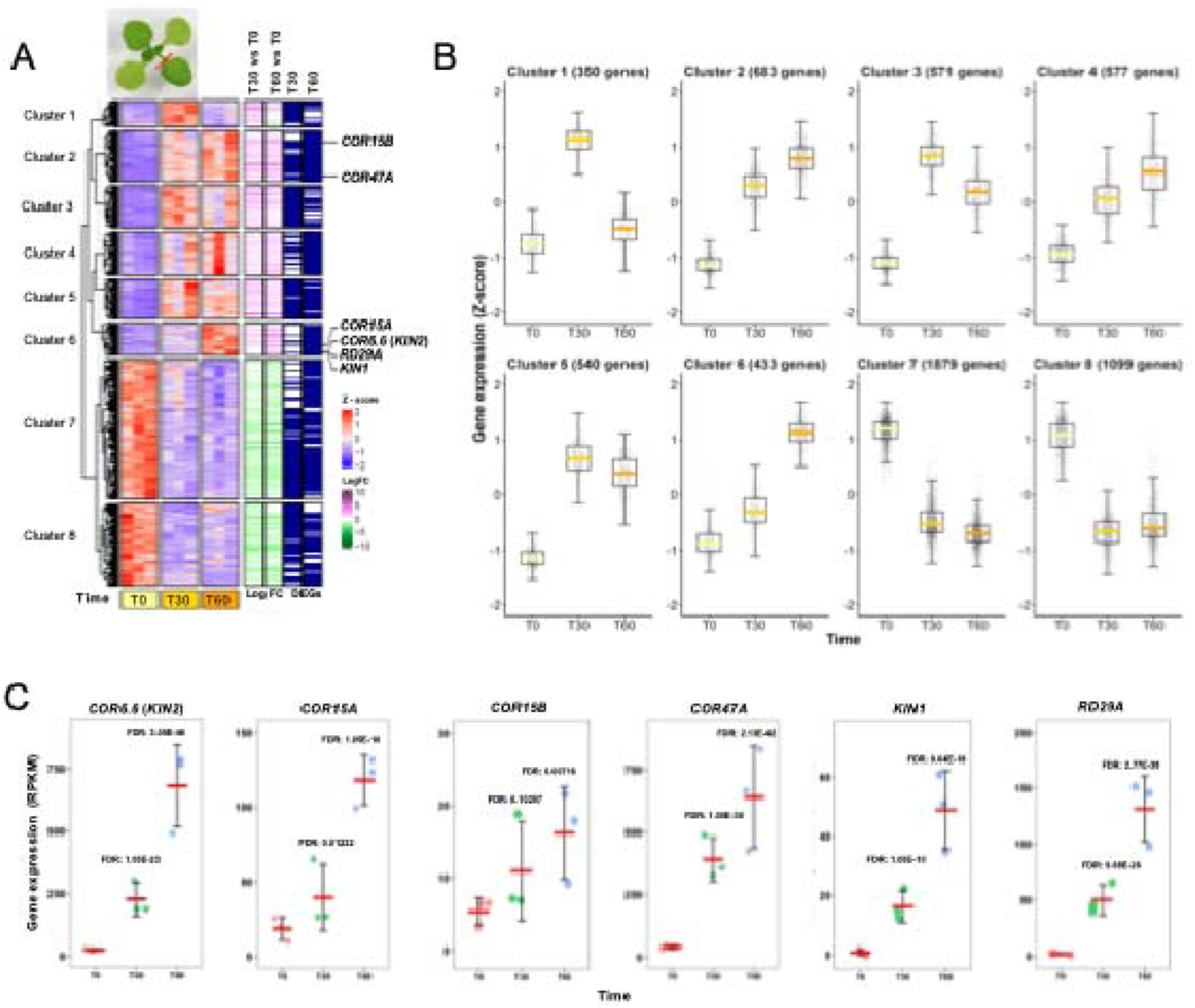
Cold-responsive genes are up-regulated in wounded tissues. A) Hierarchical clustering of the 6132 genes that responded to wounding at one or more time points. 1.5 fold-changes for the comparisons ‘T30 vs. T0’ and ‘T60 vs. T0’ are shown on the right. Statistically significant differentially expressed genes (DEG) are marked with blue stripes. Notably, cold-responsive marker genes are enriched in Cluster 2 and 6, indicating their up-regulation following wounding. B) Expression patterns of the eight gene clusters identified in the heatmap. Clusters 1–6 represent genes up-regulated in response to wounding, while Clusters 7 and 8 comprise down-regulated genes. C) Expression profiles of six cold-responsive marker genes: *COR6.6* (*KIN2*), *COR15A*, *COR15B*, *COR47A*, *KIN1*, and *RD29A*. Red bars indicate the mean, and error bars represent standard deviation. FDR (False Discovery Rate) values are indicated for each comparison.

### Wound sites had low temperature compared to leaf blade

Given the activation of cold-responsive genes in wounded tissues, we directly measured the temperature of a wounded leaf to determine if there was a causal reduction in temperature after wounding. We cut the tip of *Arabidopsis* leaves grown under a short-day condition in soil. The real-time temperature profile was monitored using an infra-red, thermal imaging camera (Figure 2A). Consistent with the gene expression results (Figure 1), infra-red thermal imaging revealed an immediate temperature reduction (within seconds) at the cut sites (Figure 2B, Supplementary Video 1). On average, the local cooling at 5 minutes following wounding was ∼ 1.5 °C lower than the rest of the leaf (Figure 2B). We also observed similar immediate temperature reductions following wounding in *Solanum lycopersicum* (*Sl*), *N. benthamina* (*Nb*), *Epipremnum aureum* (*Ea*) and *Kalanchoe daigremontiana* (*Kd*) (Figure 2C). In addition to these room temperature experiments (22°C), wound-induced cooling was also observed when the environmental temperature was at 4°C and 40°C (Supplementary Figure 2). To confirm the cooling effect at wound sites was not due to thermal imaging error, we used a fine-tip thermocouple to measure temperature at wound sites (Figure 2D) and found that the hand measured thermocouple temperatures also displayed an average of ∼ 1.5 °C lower at the wound site (Figure 2E). Thus, two independent methods confirmed that an immediate cooling occurs at the wound site on a leaf, which coincided with the activation of cold-responsive genes (Figure 1).

**Figure 2:**
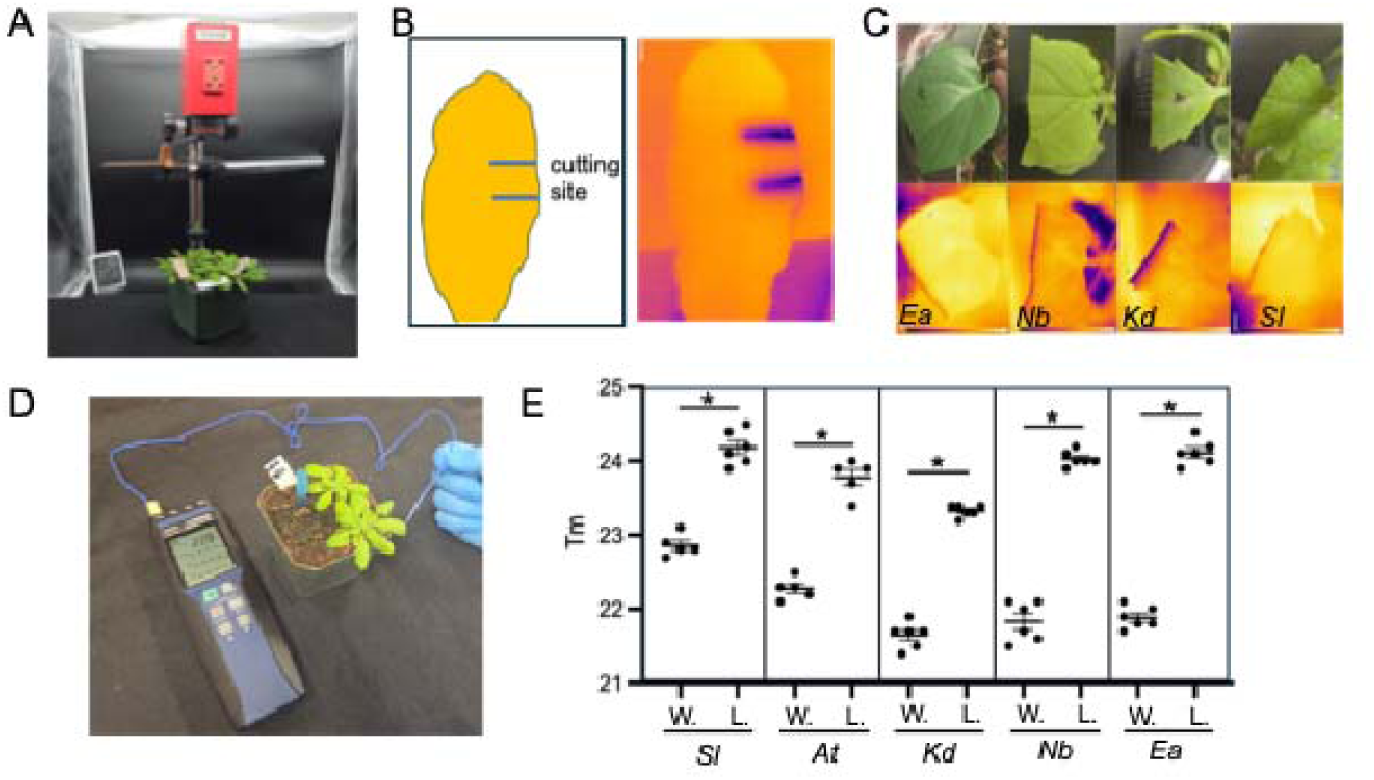
Localized temperature reduction at leaf wounding sites. A) Arabidopsis plants were imaged in a photobooth using Fotric’s 616C R&D thermal camera. B) Within 2-5 seconds after mechanical wounding, cutting sites exhibited a temperature reduction of 1.5-2.0°C, as indicated by the dark purple hue of the wounding sites in the thermal image. C) Temperature reduction at cutting sites were observed in *Epipremnum aureum* (*Ea*), *Nicotiana benthamiana* (*Nb*), *Kalanchoe daigremontiana* (*Kd*), and *Solanum lycopersicum* (*Sl*). Upper panels: RGB images; Lower panels: thermal images. D) Measuring wounding site and leaf blade temperature using a fine-tip thermocouple. E) Reduced temperature at wound sites were detected in various plants using a thermocouple. *:p<0.05 using unpaired t-test with a two-tailed hypothesis. W: wound; L: leaf. Each dot represents a measurement from one leaf.

Since the temperature reduction was only observed at the cutting site (∼ within 1-2 mm of the wound), we further monitored the spatiotemporal pattern of a cold responsive promoter (*proCOR15A::GUS*) after wounding (Baker *et al*., 1994). *COR15A* is induced by freezing (Steponkus *et al*., 1998) and low ambient temperatures (Franklin & Whitelam, 2007). Notably, it was also among the wound-induced genes in our RNA-seq experiment (Figure 1A and 1C). We observed an induction of *COR15A* promoter activity at as early as 10 mins after cutting, and the staining signal was preferentially detected at the cutting site (Figure 3A). The localized expression pattern became evident after 1 hour and was maintained in an upregulated state for 2-3 days (Figure 3A). In contrast, an auxin-responsive promoter (*proDR5::GUS*) and an indicator of cell division (*proCYCLINB1;1::GUS*) were not activated at the cutting site of a leaf residue within the 7-day period we monitored (Figure 3A), indicating that limited auxin response and cell proliferation occurred at these wound sites. Our data showed that wound-induced cooling coincided with the temporospatial expression pattern of cold-responsive genes.

**Figure 3:**
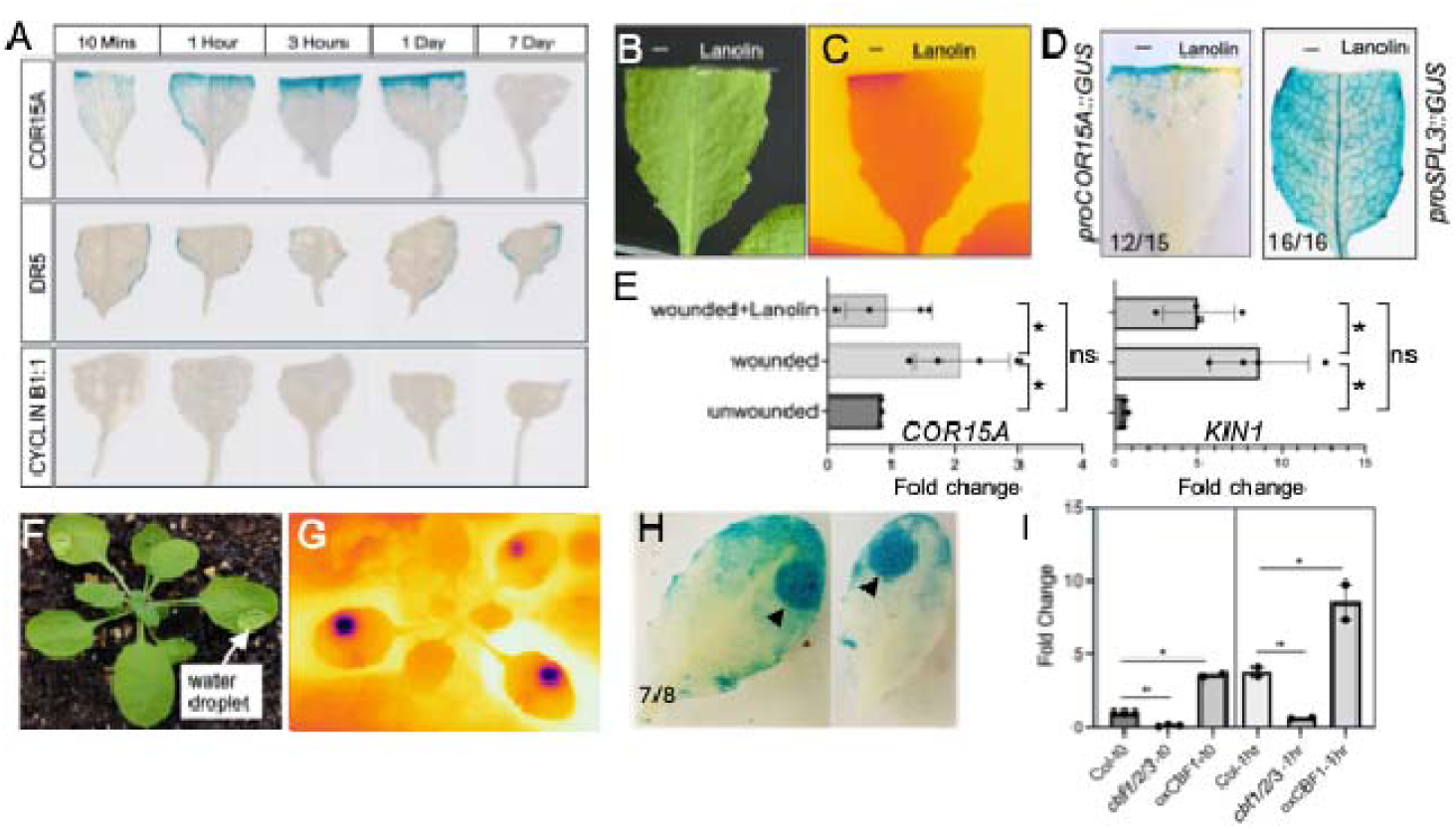
Evaporative cooling activated *COR15A.* A) Expression pattern of *COR15A*, *DR5* and *CYCLIN B1;1* promoters after cutting. Staining of GUS reporters driven by *proCOR15A* (top), *proDR5* (middle) and *proCYCLIN B1;1* (bottom) after wounding. *proCOR15A::GUS* was induced as early as 1 hour after cutting. *proDR5* and *proCYCLIN B1;1* were not induced at wound sites on leaf residue within 7DAC. B) and C) RGB (B) and thermal (C) images showing a wounded leaf with lanolin applied on the right side. Note the loss of cooling on the lanolin applied side. D) Staining of wounded leaves carrying *proCOR15A::GUS* (left) and *proSPL3::GUS* (right). Lanolin was applied on the right cutting edge of each leaf. E) qPCR of *COR15A* and *KIN1* in wounded tissues with or without lanolin. Bars represent standard error. *: p<0.05 using an unpaired (Welch’s) t-test with a one-tailed hypothesis. Each dot represents a biological replicate. F) and G) RGB (F) and thermal (G) images showing leaves with water droplets. A reduced temperature was associated with water droplets. H) Staining of *proCOR15A::GUS* in leaves with water droplets. Arrowheads indicate the location of water droplets. of x/y: x leaves showed observed pattern in a total of y leaves. I) Wound-induced activation of *COR15A* depended on CBFs. Wound samples were harvested at 1 hour or 1 DAC. Bars represent standard deviation. *: p<0.05 using an unpaired (Welch’s) t-test with a one-tailed hypothesis. Each dot represents a biological replicate.

To test a hypothesis that the induction of cold-responsive gene was due to evaporative cooling, we applied lanolin, a waxy water-repellent substance, onto half of a wound site to selectively block water evaporation (Figure 3B). We observed that the region with lanolin had a higher temperature than the exposed side of wound (Figure 3C). Consistently, the induction of *COR15A* promoter was compromised at lanolin-treated half (Figure 3D). As a control for lanolin treatment, *proSPL3:GUS*, a gene unresponsive to cold, did not show a reduction in the region applied with lanolin (Figure 3D). Using qPCR, we also observed that wound induction of *COR15A* and *KIN1* transcripts was restored to basal level in the presence of lanolin (Figure 3E). As a complementary approach, we generated wounds on a seedling submerged under water (Supplementary Figure 3). After two hours, the seedlings were stained for the pattern of *proCOR15A::GUS* (Supplementary Figure 3). Compared to control plants with leaves cut in the open air, the submerged plants did not show activation of *COR15A* promoter at its cutting sites (Supplementary Figure 3A and B). These results suggest that wound associated evaporative cooling may contribute to the transcriptional activation of cold responsive genes.

*COR15A* are often shown to be induced under freezing stress (Steponkus *et al*., 1998; Franklin & Whitelam, 2007). Compared with freezing, evaporative cooling only mildly reduced leaf temperature in our studies (Figure 2). To determine whether evaporative cooling was sufficient to induce *COR15A* promoter activity, we applied a drop of room-temperature water on leaf surface for several hours and monitored *proCOR15A::GUS* activity (Figure 3F, 3G). We observed a low temperature associated with the site of water droplet and a preferential activation of *proCOR15A::GUS* underneath the droplet (Figure 3H). Since no wounding was involved in this assay, the results suggest that a persistent evaporative cooling is sufficient to locally induce a subset of cold-responsive gene expression.

Next, we explored the genetic requirement of CBF transcription factors in wound-induced activation of *COR15A*. We compared the mRNA level of *COR15A* at the wound site from Col-0 and a triple mutant of *CBFs*, *cbf1/2/3* (CS73345) (Xie *et al*., 2021), and a transgenic line overexpressing *CBF1* (*35S::CBF1*, CS69498) (Gilmour *et al*., 2004). Using qPCR, we observed an induction of *COR15A* in Col-0 after wounding, consistent with the RNA-seq results and the GUS reporter (Figure 3I). Such induction was abolished in the *cbf1/2/3* triple mutant, implicating a CBF-dependent mechanism (Figure 3I). *COR15A* level was generally higher in the CBF1-overexpression line even in the absence of wounding (Figure 3I). These observations indicate that CBFs contribute to the wound-associated activation of *COR15A*.

### A deep learning-based workflow to monitor wound temperature

Thermal imaging of evaporative cooling at the wound site provides a real-time, non-invasive approach to monitor wound healing (Figure 2). With an assumption that healed tissue would restore wound site temperature due to the gradual loss of evaporative cooling (Figure 4A), we proposed a DL-based workflow to measure the temperatures of the wound and blade sites (Figure 4B). RGB and thermal images were first registered based on the corner points of the checkboards (Figure 4C). Next, a DL-based segmentation model, YOLO-seg, was employed to segment the blade and wound sites in RGB images (Figure 4D). The resulting segmentation masks were then mapped onto the raw thermal images to extract the temperatures of the blade and wound areas (Figure 4E). Finally, the temperature differences were calculated based on the extracted temperatures.

**Figure 4:**
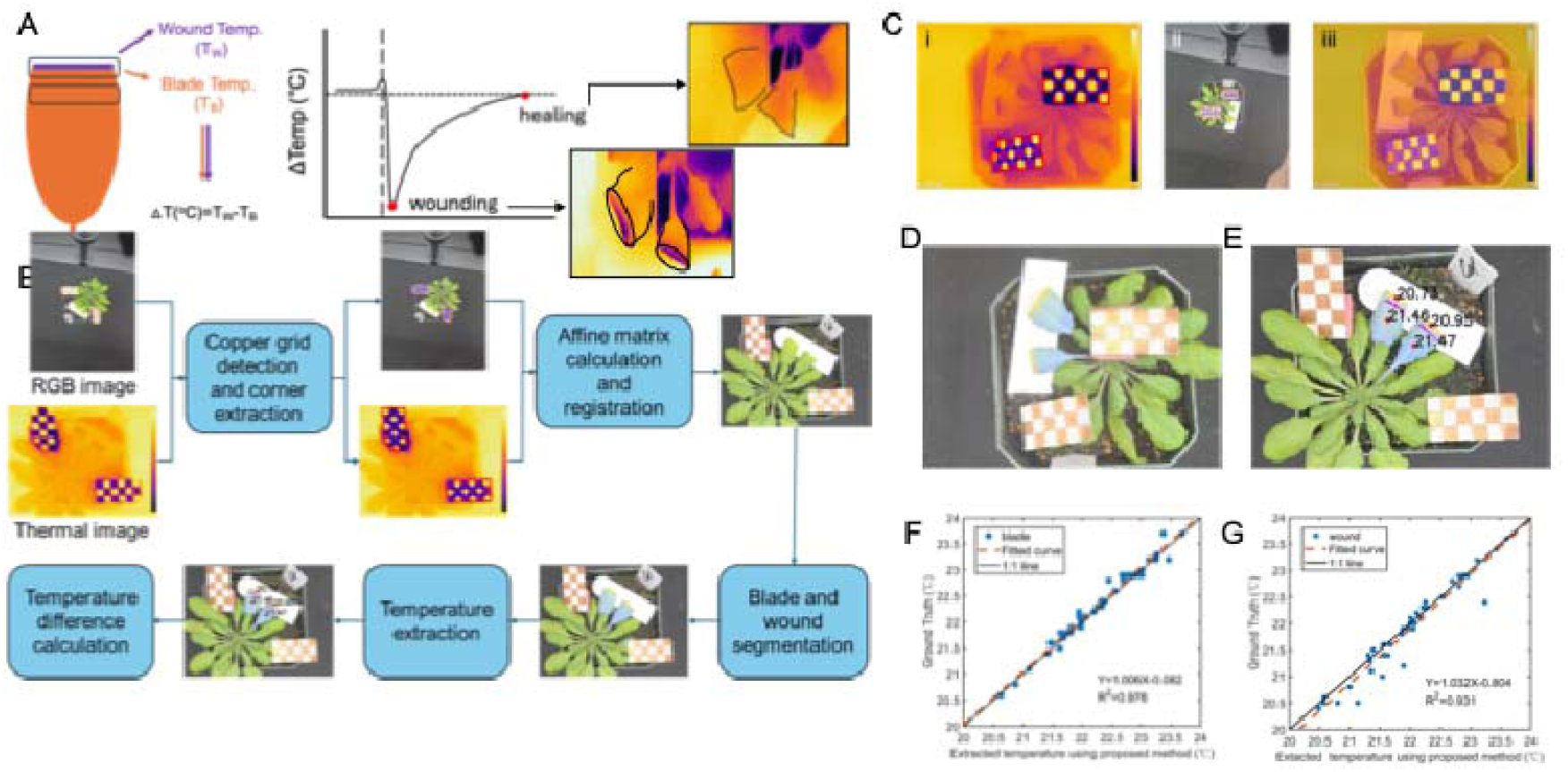
DL-based workflow for temperature difference measurement between blade and wound extraction. A) A diagram showing the method to measure temperature difference between wound site and leaf blade. Representative images of leaves at 30 mins and 3 days after wounding. Leaf margins were outlined. Note the lack of evaporative cooling in leaves at 3 DAC. B) The overview of the DL-based method for temperature difference extraction. C) Thermal and RGB image registration. (i-ii): Copper grid detection (red bounding boxes) and corner extraction (blue dots) results on both RGB and thermal images, respectively. (iii): The aligned results by overlaying the transformed RGB image onto the thermal image. D) Segmentation results of the blade (blue masks) and wound (yellow masks). E) Temperature extraction of the blade and wound based on the segmentation results. F) Linear regression between DL-extracted temperatures and the ground truth of blade sites. G) Linear regression between DL-extracted temperatures and the ground truth of wound sites.

The performance of the proposed method on the testing datasets of 22 images demonstrated high accuracy in detecting copper grids of the checkerboards for both RGB and thermal images. Specifically, the Precision, Recall, mAP@0.5, and mAP@0.5:0.95 were 99.9%, 99.6%, 99.5%, and 97.3%, respectively, for RGB images, and 99.9%, 100.0%, 99.5%, and 97.0% for thermal images. The segmentation performance for the leaf blade and wound sites on RGB images was also strong, with overall Precision, Recall, mAP@0.5, and mAP@0.5:0.95 reaching 97.3%, 96.7%, 98.0%, and 70.9%, respectively. Furthermore, the extracted temperatures for the leaf blade and wound sites using the proposed method closely correlated with ground truth measurements obtained manually from thermal images and the thermocouple measurements (Figure 4F and 4G). These results indicated that the proposed DL-based workflow accurately estimated temperatures, with leaf blade predictions showing slightly higher accuracy than those for wound sites. Specifically, the mean absolute errors (MAE) were 0.09°C for blade sites and 0.15°C for wound sites, with R² values of 0.976 and 0.931, respectively. In summary, the DL-based workflow can automatically collect both leaf blade and wound site temperatures using RGB and thermal imagery.

### CBFs were required for normal wound healing

We then used the DL-based workflow to monitor the temperature differences between wound site and leaf blade over a 4-day period in wild type Col-0 and the triple mutant of *cbf1/2/3* (Xie *et al*., 2021). We defined “ΔTm” as the difference between wound and leaf blade temperature and expected it to approach zero upon wound sealing (Figure 4A). In Col-0, the cooling effect was readily detected immediately after cutting with a temperature difference around 1.5°C (Figure 5A). The temperature difference between the wound site and leaf blade decreased between one and three days after cutting (DAC) (Figure 5A), but by 4 DAC no temperature difference was detected, indicating that the wound was healed (Figure 5A). At both 2 and 3 DAC, the *cbf1/2/3* mutant showed a greater temperature difference between the leaf blade and wound site than Col-0, suggesting a delayed healing process (Figure 5A and 5B, Supplementary Figure 5). We also used lignin deposition as a proxy for wound healing (Xu *et al*., 2024) and observed compromised lignin deposition in the *cbf1/2/3* mutants, indicating that the CBF transcription factors are required for wound-induced lignin deposition (Figure 5D and 5E). Together, these results suggested that the CBFs contribute to wound healing, probably by sensing the evaporative cooling signal.

**Figure 5:**
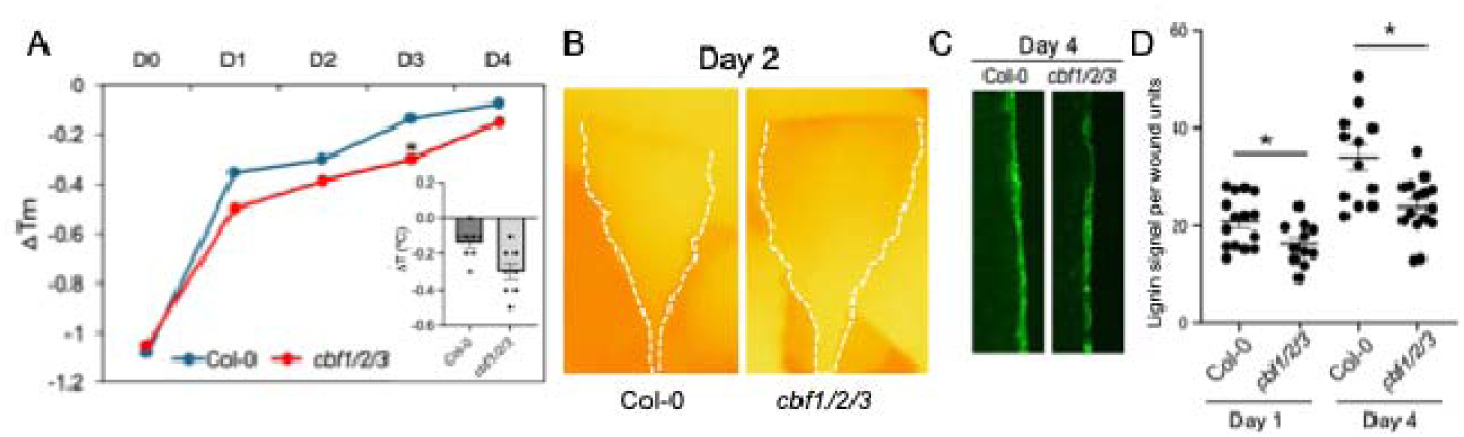
CBFs are required for normal wound healing. A) Healing curve of Col-0 and *cbf1/2/3* in 4 DAC. *cbf1/2/3* mutant showed a delayed healing process. The insertion shows temperature differences at 3 DAC in Col-0 and *cbf1/2/3*. *: p<0.05 using unpaired t-test with a two-tailed hypothesis. B) Representative thermal images from Col-0 and *cbf1/2/3* at 1 DAC and 4 DAC. Note the reduced temperature at cutting sites at 2DAC. Dashed lines indicate leaf margin. C) Representative images of lignin deposition at wound site at 1 DAC. Note the reduced lignin deposition in *cbf1/2/3* mutant. D) Quantification of lignin deposition at wound sites in Col-0 and *cbf1/2/3*. *: p<0.05 using unpaired t-test with a two-tailed hypothesis. Each dot represents a measurement from one wounded leaf.

### SA responses were compromised in *cbf1/2/3* after wounding

To further explore the downstream factors connecting CBFs to wound healing, we compared wound-induced transcriptomes between Col-0 and the *cbf1/2/3* mutant at 0, 10 min, 60 min, and 24 h after wounding. Because cooling occurs only at the wound sites, we harvested tissues within approximately 5 mm of the cutting site (Figure 6A and Supplementary Figure 6A). At each time point, we detected 99, 46, 186, and 151 differentially expressed genes (DEGs) between Col-0 and *cbf1/2/3*, respectively (Supplementary Figure 6B). We then clustered the 335 DEGs that were differentially expressed between Col-0 and *cbf1/2/3* in at least one time point (Figure 6A). As expected, we observed the activation of cold response following wounding (Figure 6A and 6B). Notably, CBFs were rapidly activated within 10 min after wounding, and their expression returned to steady-state levels within 1 h (cluster 1) (Supplementary Figure 6C). This transient activation preceded a wave of CBF-activated genes observed at 60 min in cluster 7 (Figure 6A). In the *cbf1/2/3* mutants, genes identified as CBF transcriptional targets, based on published ChIP-seq data (Song *et al*., 2021), were overrepresented among the downregulated DEGs at 10 min, further supporting that these genes are regulated by CBFs upon wounding (Supplementary Figure 6D).

**Figure 6.**
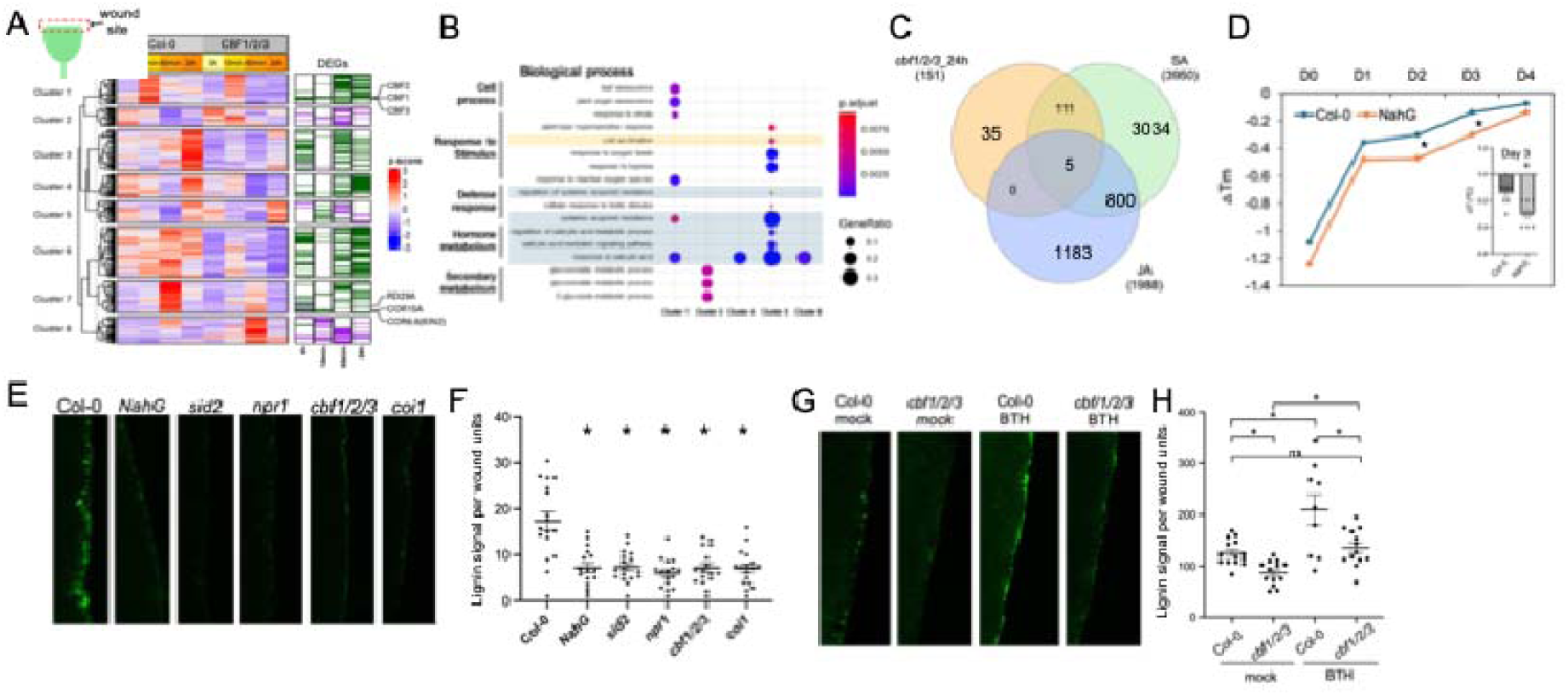
CBF-mediated regulation of wound-induced SA response and tissue healing. A) Heatmap showing hierarchical clustering of 335 DEGs between Col-0 and the *cbf1/2/3* triple mutant following wounding. CBF1, CBF2, and CBF3 were rapidly induced within 10 min after wounding, followed by activation of cold-responsive genes after 1 hour. B) Gene Ontology (GO) enrichment analysis of DEGs reveals significant overrepresentation of salicylic acid (SA)-responsive pathways. C) Venn diagram illustrating the overlap between SA-, JA-, and CBF-regulated genes at 24 hrs after wounding. Among CBF-dependent DEGs at 24 hr, 111 genes overlap with SA-responsive genes, whereas no overlap is observed with JA-responsive genes. SA and JA responsive genes were defined as in (Yang *et al*., 2017) D) Wound healing kinetics in the *NahG* mutant, monitored by thermal imaging. The temperature difference (ΔT) between wounded and unwounded tissue was significantly greater at 2 and 3 days post-cutting compared to Col-0. E) Representative images of lignin deposition at wound sites in Col-0 and SA-related mutants. *Sid2*, *NahG* and *npr1* exhibited reduced lignin accumulation compared to Col-0. Lignin deposition was detected by its autofluorescence under UV excitation using fluorescence microscopy. Coi1 was used as a positive control (Xu *et al*., 2024). F) quantification of lignin deposition for Col-0 and mutants showed in E. *: p<0.05 using unpaired t-test with a two-tailed hypothesis. G) Representative images of lignin deposition at wound sites in Col-0 and *cbf1/2/3* with mock or BTH treatment. H) Quantification of lignin deposition in G. *: p<0.05 using unpaired t-test with a two-tailed hypothesis.. Ns: not significant. Each dot represents a measurement from one wounded leaf.

As part of wound response, the SA response was activated following wounding and inhibited wound-induced regeneration (Hernández-Coronado *et al*., 2022; Tran *et al*., 2023). Noteworthy, gene ontology (GO) analysis of CBF-dependent DEGs revealed significant enrichment of salicylic acid (SA)–related processes, including response to SA, systemic acquired resistance (SAR), and SA signaling pathways (Figure 6B). These GO terms were enriched in clusters 1, 4, 5, and 8, which include genes that were failed to be activated in the *cbf1/2/3* mutant upon wounding (Figure 6B). These results indicate that CBFs are required for the wound-induced SA response. Notably, at 24 hours after wounding, 111 of 151 CBF-regulated genes overlapped with the SA regulon (Yang *et al*., 2017). In contrast, although activation of JA and ABA pathways are important for wound repair (Koo & Howe, 2009; Xu *et al*., 2024), among the 335 CBF-regulated genes, only 22 and 30 overlapped with previously reported ABA- and JA-responsive genes, respectively (Wang *et al*., 2011; Yang *et al*., 2017). Wound-induced activation of the *LOX2* promoter, a JA-responsive marker, was not blocked by lanolin treatment (Supplementary Figure 6E), suggesting that CBFs may not be essential for wound-induced JA activation. . At 24 hours after wounding, none of the 151 DEGs overlapped with JA-responsive genes (Figure 6C) (Yang *et al*., 2017), further confirming that the CBFs have limited influence on wound-induced JA response within this time frame. On the other hand, *COR15A* induction was not altered in mutants defective in ABA biosynthesis (*aba3-1*) (Supplementary Figure 6F), so wound-induced activation of *COR15A* is likely independent of JA and ABA signaling. These results strongly support that CBFs regulate a wound-induced SA regulon. We therefore tested whether blocking SA pathway affects wound healing. In *NahG* transgenic plants, which exhibit reduced SA accumulation due to overexpression of a *Pseudomonas*-derived salicylate hydroxylase, wound healing was delayed, as indicated by a larger temperature difference between the wound site and surrounding leaf tissue at 2 days after cutting (Figure 6D). Furthermore, wound-induced lignin deposition was decreased *NahG*, *sid2* (mutant defective in Isochorismate Synthase 1-controlled SA biosynthesis), and *npr1* (an SA receptor), as observed in *cbf1/2/3* and *coi1* (Figure 6E and 6F). To test the capacity of exogenous SA to rescue wound-induced lignin deposition in the *cbf1/2/3* mutant, we pre-treated Col-0 and *cbf1/2/3* plants with BTH, an SA analog, 24 hours prior to cutting. At 1-day post-cutting, BTH treatment significantly enhanced lignin deposition at the cutting site in mutant plants and restored it to the level in wild type (Figure 6G and 6H), suggesting that SA may promotes lignin deposition downstream of CBF function. Together with the gene expression profile, we conclude that SA response acts downstream of CBFs to promote wound repair.

## Discussion

The healing of wounded plant tissues is an essential process that protects plants from subsequent damage caused by opportunistic microbial invaders, herbivorous insects, and pathogenic organisms. Despite its importance, a major challenge in understanding wound healing lies in defining the point at which a wound can be considered fully healed. We provide experimental evidence that wound-induced evaporative cooling, driven by abrupt yet subtle temperature changes, can trigger the activation of cold response pathways. Thermal imaging and the deep learning workflow provided a reliable and sensitive method for us to track wound healing through the gradual increase in wound site temperatures. Presumably, when the temperature at the wound site is indistinguishable from the remaining undamaged leaf blade, the wound is sealed. It is possible that the wound is not entirely healed at this point because the sensitivity of the camera may limit our ability to measure relatively small differences in temperature loss. Regardless, thermal imaging combined with molecular changes (e.g. induction of *COR15A* and lignin deposition) offers an important method to evaluate physical and molecular responses related to the healing end point. This is a key step towards a more refined understanding of the plant wound healing responses.

In this study, we found that evaporative cooling following injury was sufficient to activate a cold response dependent on CBF transcription factors. Loss of *CBFs* resulted in deficient lignin deposition at the wound site and impaired wound healing, highlighting the biological importance of seemingly slight temperature reductions. The observation that *COR15A* was induced by evaporative cooling from a persistent water droplet further support that plants are highly sensitive to temperature changes across different temporal scales. While many studies on thermal sensing have focused on plant responses to physiologically extreme temperatures, such as freezing and heat stress, ambient temperature fluctuations also profoundly impact development and stress responses (Dong *et al*., 2020; Kerbler & Wigge, 2023). At the cellular level, molecular thermosensors respond quantitatively to fine temperature variations. For instance, in *Arabidopsis*, hypocotyl elongation is regulated by ambient temperatures ranging from 17°C to 37°C (Hahm *et al*., 2020; Qiu *et al*., 2021). The number of PhyB-containing nuclear bodies gradually decreases as the temperature rises from 12°C to 27°C, with a small 4°C difference being sufficient to induce PhyB dissociation and hypocotyl elongation (Qiu *et al*., 2019). These responses to subtle temperature difference is similar to our observations that sudden evaporative cooling of ∼ 1.5 °C at wound site appears sufficient to elicit a physiological response.

Several lines of evidence indicate connections between low temperatures and callose or lignin accumulation (Schütte *et al*., 2025). In pepper, two lignin biosynthesis genes, *CaCAD1* and *CaPOA1*, are induced by prolonged cold exposure (Xiao *et al*., 2025). Our results showed that CBF promotes the wound-induced SA response. It is therefore plausible that SA serves as a link between sensing evaporative cooling and the deposition of callose and lignin, given its established role in pathogen-induced callose accumulation (Tateda *et al*., 2014) and its crosstalk with lignin biosynthesis (Delaney *et al*., 1994; Wang *et al*., 2025). Notably, DEGs between Col-0 and *cbf1/2/3* mutants at 1 h and 24 h after cutting are not enriched for CBF-binding motifs, suggesting that CBFs likely regulate SA-responsive genes indirectly through yet unidentified intermediates. Interestingly, previous studies have shown that *cbf1/2/3* mutants did not exhibit altered activation of SA-responsive genes (*PR1* and *PR5*) in response to bacterial infection or exogenous SA treatment, indicating that CBFs are not required for pathogen-induced SA signaling (Li *et al*., 2020; Li *et al*., 2024). We also assessed bacterial pathogen growth and the induction of *PR1* and *PR5* following BTH treatment in our *cbf1/2/3* mutant and observed no significant differences compared to the wild type (Supplementary Figure 7). Together, these findings suggest that additional wound-associated factors that are not present during pathogen infection, may act synergistically with CBFs to activate wound-induced SA signaling. Overall, these data support the existence of an intermediate regulatory component linking CBF transcription factors to the wound-induced SA response.

The deep learning approach developed in our study combined multiple imaging modalities, i.e., RGB and thermal images, to detect the wound sites with precision. This integrated approach not only identified wounds but also quantified localized temperature variations—a key indicator of physiological stress, disease progression, and healing efficiency (Wen *et al*., 2023). By automating the analysis of these multimodal datasets, the method significantly reduces reliance on labor-intensive manual assessments, reduces observer bias and enables high-throughput phenotyping. The workflow’s capability to track thermal profiles at specific injury sites offers deeper insights into plant resilience mechanisms. Overall, these capabilities can accelerate fundamental research, improve crop management strategies, and enhance data-driven decisions to boost agriculture productivity and long-term sustainability.

## Materials and methods

### Plant materials and growth conditions

*Arabidopsis* wild type and mutants used in this study were summarized in supplementary Table 2. Seeds were sown on Sungro Propagation Mix Question soil and placed in 4°C cold room for 3 days prior to being transferred to a short-day growth room with 23°C/19°C day/night temperatures and 45% humidity. Plants were watered weekly and placed in a random block design to reduce the impact of environmental factors. Plants for thermal imaging and GUS staining were 4-6 weeks old.

### GUS staining

Leaf tissues were submerged in a GUS staining solution [100 mM sodium phosphate (pH 7.0), 1 mM EDTA (pH 8), 1% Triton-X-100, 5 mM potassium ferrocyanide, 5 mM potassium ferricyanide and 1 mg/ml X-Gluc] and vacuum infiltrated for 10 min. Following incubation at 37°C (3 hours for proLOX2:GUS, 5 hours for proCOR15A:GUS, proDR5:GUS, and proCYCB2:GUS. The staining solution was then replaced with 70% ethanol and washed several times until the tissue was clear. Once cleared, the samples were blotted dry, placed face-down on a CanoScan LiDE 400 PDF scanner, and scanned at 1200 dpi with all color correction and blemish removal settings disabled.

### qPCR

RNA extraction was performed using TRIzol reagent (Invitrogen). The qPCR was performed in the Applied Biosystems QuantStudio 1 Real-Time PCR system with SYBR Green master mix (Applied Biosystems). PCR conditions were set as follows: 95°C for 5 mins, 40 cycles of 95°C for 15s, 56°C for 30s and 72°C for 20-30s. *TUBULIN 2* (*AT5G62690*) was used as a reference gene. We confirmed that *TUBULIN 2* was not a wound-induced DEG. The delta-delta CT was used for fold-change calculation after confirming the PCR efficiency was at least more than 95%. The oligonucleotides used are COR15A-F: 5’-AACTCAGTTCGTCGTCGTTTCTC-3’; COR15A-R: 5’-CCCAATGTATCTGCGGTTTCAC-3’; TUB-F: 5’-AGCAATACCAAGATGCAACTGCG-3’ and TUB-FR: 5’-TAACTAAATTATTCTCAGTACTCTTCC-3’; PR1-F: 5’GCCTTCTCGCTAACCCACAT-3’ PR1-R: 5’-CGGAGCTACGCAGAACAACT-3’; PR5-F: 5’-ACTGTGGCGGTCTAAGATGTA-3’ PR5-R: 5’-CGTATTTGCAATCTCCCGATCC-3’ KIN1-F: 5’-GCTGGAGCTTCCGCGCA-3’ KIN1-R: 5’-TTATAACTCCCAAAGTTGACTCGG-3’

### Acquisition of thermal imaging

Fotric’s 616C R&D station equipped with a 40.5 mm thermal lens and AnalyzIR software was used to acquire thermal images (FOTRIC). The station was situated in NEEWER’s 20”x20”x20” Studio Light Box with LED lights strung in the canopy and black fabric lining the bottom of the booth. The height of the camera mount was adjusted to create a 15 cm object distance and emissivity is designated as 1. A 3x6 cm^2^ checkboard with alternating copper and paper cells was used as an RGB image alignment marker. The checkboard was mounted on a 30° rubber wedge to avoid directing the heat signature from the camera body back towards the camera.

Arabidopsis seeds were grown between four and six weeks prior to cutting. On the day of cutting, half of the leaf blade was removed with ceramic scissors. A plastic stake was placed below the cutting site with forceps to minimize soil temperature affecting image quality. At 30 minutes post-cutting, the plant was transferred to the photo booth. Two pre-chilled checkboard alignment markers were placed within the camera’s field of view and away from any leaves. After adjusting the focus until the cutting site was well defined, snapshots were taken with the AnalyzIR workstation. A report of experimental parameters and a thermal data CSV file were generated for each image. Before removing the plant from the chamber, RGB images were taken using a Samsung Galaxy S24 phone mounted on a metal stand 15 cm away from pots. Color correction and image processing function was disabled.

### Manual temperature measurement from thermal images

In AnalyzIR, the average temperature from a line tracing the cutting site was recorded as the cutting site temperature. The line was then moved to the blade area of the same leaf to measure leaf blade temperature. The temperature difference between the wounding site and leaf blade was used as a quantitative readout of cooling effect. In the time-lapse experiments of 24 hrs, the thermal camera was positioned in a growth chamber in order to monitor plants in constant growth conditions. An imaging rate of 6 seconds per frame was used to capture the time lapse, and AnalyzIR segmentation and graphing functions were used to extract temperature values.

### Manual temperature measurement from thermocouple measurements

Using an Omega HH374 Data Logger Thermometer equipped with a Type K beaded probe (Omega), temperatures on the leaf blade and the cutting site were measured. All measurements with the thermocouple were performed on the same timeline as the thermal image measurements.

### Lignin deposition quantification

Cut leaf samples were transferred to glass slides and imaged with the Zeiss Axio Imager 2 (Zeiss). To capture the lignin autofluorescence, an excitation wavelength of 488 nm and a filter for 535 nm light were used. All exposure times, gamma settings, and fields of view were standardized across all samples. ImageJ was used to remove background autofluorescence and quantify the average lesion area per micron of wounding site length.

### RNA-seq analysis

For the RNA-seq gene expression analysis following wounding, total RNA was extracted from two-week-old Arabidopsis (Col-0 ecotype) leaf samples subjected to wounding stress. Samples were harvested at three time points: prior to treatment (time 0), 30 minutes post-wounding, and 60 minutes post-wounding. The experiment included three biological replicates per condition. For the RNA-seq experiment comparing wound responses in Col-0 and in the *cbf1/2/3* mutant, tissues within 1 cm of wounding sites were harvested following cutting off 1/3 of a leaf using scissors.

Libraries were prepared using the TruSeq RNA library prep kit (Illumina) according to the manufacturer’s protocol and sequenced with the Illumina Nextseq platform. Read quality was assessed using FastQC version 0.11.8 (Babraham Bioinformatics, Cambridge, UK), and low-quality reads and adapter sequences were removed with Trimmomatic version 0.36 (Bolger *et al*., 2014). The resulting high-quality reads were then aligned to the TAIR10 reference genome using HISAT2 version 2.2.0 (Kim *et al*., 2015), and raw read counts were quantified using the featureCounts function from the Sub-read package version 2.0.0 (Liao *et al*., 2013). Differential gene expression analysis was conducted using the edgeR package in R (Robinson *et al*., 2010). Genes with low expression levels were filtered by excluding those with fewer than 1 count per million in at least five libraries. Normalization of read counts was performed using the trimmed mean of M-values (TMM) method, implemented with the calcNormFactors function in edgeR (Robinson & Oshlack, 2010). To control for multiple testing, p-values were adjusted using the Benjamini-Hochberg procedure (Benjamini and Hochberg, 1995). Genes were considered differentially expressed if they had a false discovery rate (FDR) ≤ 0.05 and exhibited a minimum 1.5-fold change in expression. Comparisons were made between gene expression levels at 30 and 60 minutes after wounding versus unwounded samples (time 0). Gene Ontology (GO) term enrichment analysis was performed using Gene Set Enrichment Analysis (GSEA) (Yi et al. 2013) to identify biological processes associated with the differentially expressed genes. For the Col-0 vs *cbf1/2/3* mutant comparison, GO enrichment analysis was performed using the compareCluster function in clusterProfiler (Yu *et al*., 2012).Hierarchical clustering was performed using the ComplexHeatmap package in R (Gu *et al*., 2016), based on the Euclidean distance and the complete-linkage method. Gene expression values in TPM (transcript per million) were normalized to z-scores for clustering analyses. Raw RNA-seq reads and the count matrix generated in this study are available at the NCBI Gene Expression Omnibus under the accession number GSE293328. Additionally, public gene expression datasets (Kilian *et al*., 2007; Ikeuchi *et al*., 2017; Fiorucci *et al*., 2022; Liu *et al*., 2022) were utilized to investigate whether cold-responsive marker genes were induced in independent experiments involving wounded *Arabidopsis* tissues, using a threshold of |log2 fold change| ≥ 1 and FDR ≤ 0.05.

### Method for testing binding sites enrichments

Enrichment analysis was performed using a hypergeometric test (phyper function in R) to determine whether genes identified as targets of CBF transcription factors, based on published ChIP-seq data (Song *et al*., 2021), were overrepresented among the differentially expressed genes in our RNA-seq experiment (DEGs). The set of expressed genes was used as the background universe and p-values were adjusted for multiple testing using the Benjamini–Hochberg method.

### Development of DL-based workflow for wound detection

The DL-based workflow consists of four main steps (Figure 4B): (1) copper grid detection and corner point extraction; (2) RGB image and thermal image registration based on affine matrix; (3) blade and wound segmentation on RGB images; and (4) mapping the segmented blade and wound to aligned thermal images and temperature difference calculation.

Two checkerboards made of foam grids and copper sheets were placed around the wounded area, and an RGB camera (Samsung Galaxy S24) and a thermal camera (600 R&D, FOTRIC) were used to collect the RGB and thermal images from the top view. Due to the different resolutions (3000x4000 for RGB and 640x480 for thermal) and varying parameters such as the focal lengths of the cameras, the RGB and thermal images were not aligned. Therefore, we extracted corners on the checkerboards for alignment. As the temperatures of the foam and copper are different, the copper is lighter, and the foam is darker in the thermal images. To detect copper grids and extract the corner points, the detection model, YOLOv8-obb (Jocher *et al*., 2023) was trained to detect the copper grids with orientated bounding boxes in RGB and thermal images, respectively. A total of 71 RGB images and 38 thermal images were annotated using Roboflow (Dwyer *et al*., 2024) with oriented bounding boxes representing copper grids. Images were then divided into training and testing datasets with a ratio of 8:2. All training images were further augmented three times, including horizontal and vertical flipping, 90-degree rotations, and additional adjustments to saturation, brightness, and exposure for the RGB images. This resulted in a total of 171 RGB images and 87 thermal images for training. The detection model was trained for 100 epochs with an input resolution of 2048×2048 pixels for RGB images and 640×640 pixels for thermal images. An initial learning rate of 0.002 and an AdamW optimizer were used on the HiPerGator high-performance computing cluster with 8 AMD EPYC ROME CPU cores, one NVIDIA DGX A100 GPU node (80GB), and 32 GB of memory. After detecting all copper grids, the four corners of each grid were extracted and the distances between each corner were calculated. Then, the closest two corners were obtained, and the center point was calculated as the corner point of these two grids. A sorting algorithm that sorts the corner points from left to right and top to down to align the same corner points on RGB and thermal images. The affine matrix was calculated based on the sorted points and the affine transformation was applied to the RGB image to register with the thermal image (Figure 4C).

To detect the blade and wound sites, a segmentation model, YOLOv8-seg (Jocher *et al*., 2023), was trained. A total of 103 images were annotated using Roboflow with polygons representing blade and wound sites. Images were then divided into training and testing datasets with a ratio of 7:3. All training images were further augmented three times, including horizontal and vertical flipping, 90-degree rotation, and rotation between ±15 degrees, resulting in 216 images for training. The segmentation model was trained for 200 epochs with an input resolution of 640×640 pixels, an initial learning rate of 0.0017, and an AdamW optimizer on the HiPerGator high-performance computing cluster. After obtaining the masks of the blade and wound sites (Figure 4D), the centroids of the two closest masks were calculated. Additionally, the mask of the wound site was eroded with a kernel size of 3×3 pixels and moved along the direction formed by the centroids to the centroid of the blade with 10 pixels. Then, the mask of the wound site and the intersection over union between the moved mask and blade mask were mapped to the raw thermal data. The mean temperature values were calculated within two masks to represent the wound and blade temperatures, respectively (Figure 4E). The temperature difference was calculated by subtracting the temperature of the wound from the temperature of the blade.

### Bacterial growth assay

*Pseudomonas syringae* DC3000 strain was grown under 28°C on King’s B (KB) solid medium (40 g/L proteose, 20 g/L glycerol and 15 g/L agar). The medium contained rifamycin for selection and cycloheximide to inhibit fungi growth. Glycerol stock of the bacterial strains stored under -80°C. Bacterial stock was streaked on plate for a 2-day growth and was cultured in liquid KB medium for an additional day. After pelleting and resuspending bacteria in 10 mM MgCl_2_=:solution, the solution was diluted until the optical density at 600 nm was adjusted to 0.025 as measured with a spectrophotometer (Eppendorf BioPhotometer). The final volume of inoculating solution was further supplemented with 0.005% Silwet L-77 surfactant to ensure even inoculation. Arabidopsis plants grown in the same conditions as those grown for thermal imaging were dipped for several seconds into the inoculating solution. After inoculation, plants were covered by transparent lids for one hour. Leaf samples were harvested 48 hours post-inoculation using a corer to collect four tissue discs from different leaves per sample. Leaf samples were ground with a homogenizer (OMNI International) and serially diluted. KB plates with 10 μL of bacteria suspension per sample were grown at 28°C for two days. Colony forming units were counted manually and normalized according to inside area of the corer.

## Acknowledgements

L.Y. is supported by the National Science Foundation (IOS-2039313) and the National Institutes of Health (R35GM143067). P.J.P.L.T. is funded by the Serrapilheira Institute (grant G-1811-25705), the National Council for Scientific and Technological Development (CNPq, grant 308349/2022-9), and the São Paulo Research Foundation (FAPESP; grant 2018/24432-0). N.C.F.D. is supported by a fellowship from FAPESP (2023/10167-1). C.L. is supported by the Research Capacity Fund (Hatch Multistate) program, project award no. 7009022, from the U.S. Department of Agriculture’s National Institute of Food and Agriculture.

---

**Figure S1:**
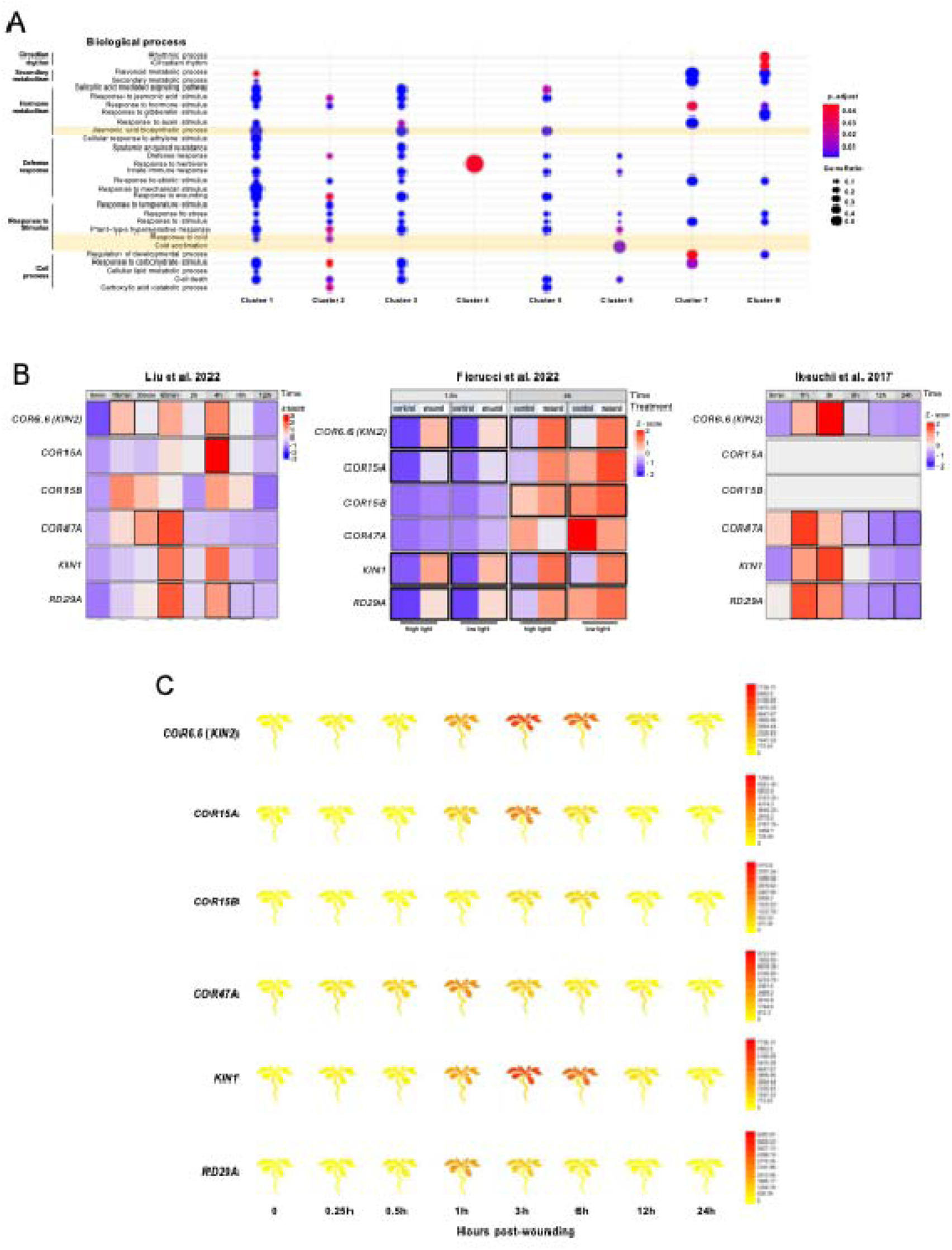
GO terms associated with wound-induced genes. A) Gene Ontology (GO) enrichment analysis indicating biological processes associated with each cluster. Each dot represents a significantly enriched GO term. Highlighted processes indicate “Response to cold”, “Cold acclimatation” and “Jasmonic acid biosynthetic process”, which are all enriched among up-regulated genes. B) Cold-responsive genes are consistently up-regulated in response to wounding across independent experiments. Gene expression is presented as z-scores, and differentially expressed genes are indicated by a thick border in each treatment. C) Up-regulation of cold-responsive genes in response to wounding. Expression of six marker genes of the cold response, represented as a cartoon-like heatmap. The image is derived from the Arabidopsis eFP Browser (Waese et al., 2017), based on AtGenExpress dataset from (Kilian *et al*., 2007). Expression levels are indicated by a gradient color scale, ranging from yellow (low expression) to red (high expression), across a time-course of 0, 0.25h, 0.5h, 1h, 3h, 6h, 12h and 24h after wounding.

**Figure S2:**
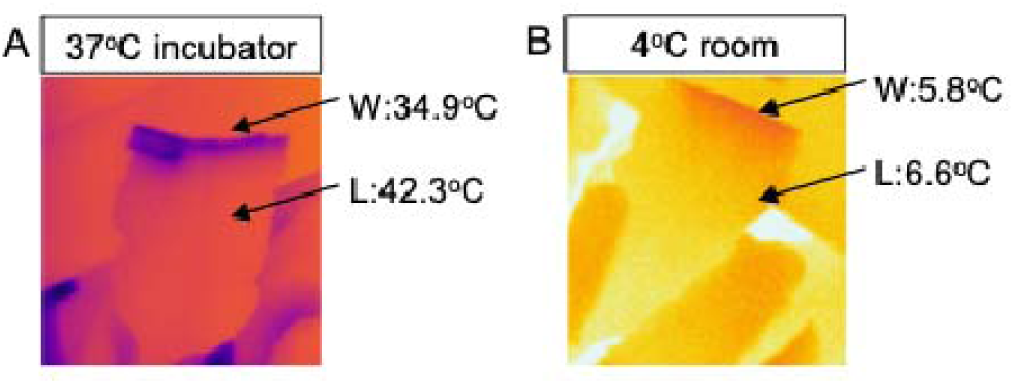
Evaporative cooling at wound site observed in cold and heat environments. Arabidopsis plants were imaged in a growth chamber at 37°C (A) or in a cold room at 6°C (B). Temperature differences between wound sites and leaf blades were observed under both conditions, with a greater difference detected in leaves at 37°C.

**Video S1: Temperature reduction occurred at wound sites immediately after cutting.**

**Figure s3:**
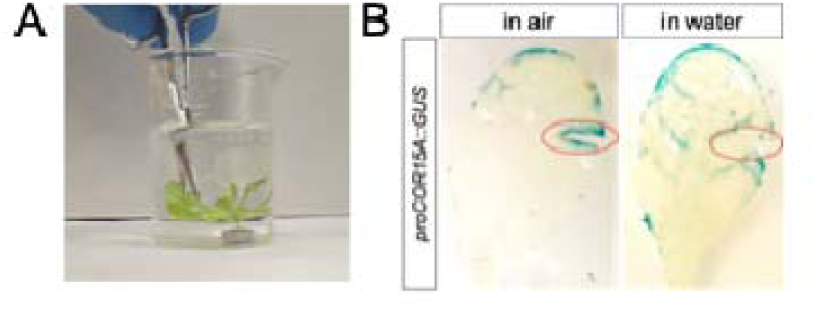
Wound-induced COR15A activation was not observed when plants were cut under water. A) An image showing a leaf being cut in water. B) Staining of *proCOR15A::GUS* in leaves wounded under water (right) or in air (left). The red circles indicate cutting sites.

**Figure S5:**
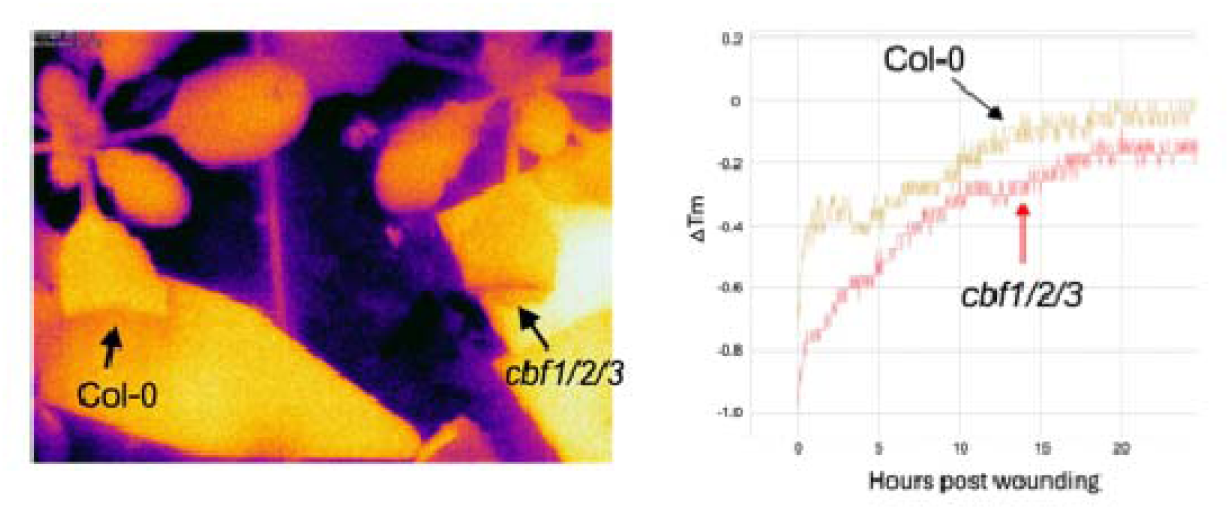
Healing curve of Col-0 and *cbf1/2/3* in the same frame. A) Thermal image snapshot of Col-0 and *cbf1/2/3* at 2 Days after cutting. B) Time course recording of temperature difference (Wound temperature – blade temperature) from a wounded Col-0 and *cbf1/2/3* leaf.

**Figure s6.**
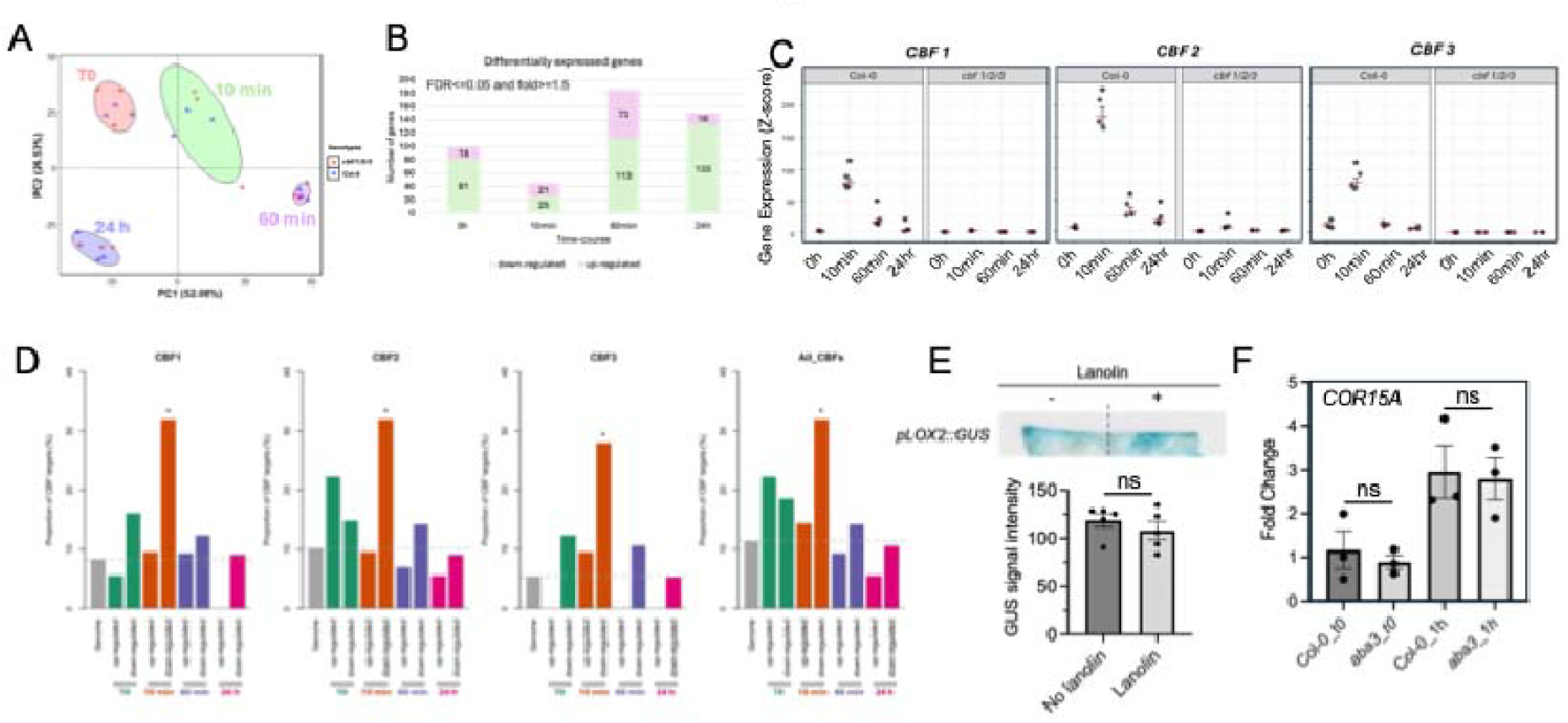
Temporal transcriptomic dynamics in response to wounding. A) Principal component analysis (PCA) of RNA-seq samples showing that time post-wounding is the primary factor driving variation in gene expression across samples. B) Number of differentially expressed genes (DEGs) identified at each time point following wounding. C) Expression dynamics of *CBF1*, *CBF2*, and *CBF3*. D) Enrichment analysis of CBF-binding motifs among DEGs at each time point. CBF-binding sites were defined based on data from (Song *et al*., 2021), and enrichment significance was assessed using a hypergeometric test, with p-values adjusted by the Benjamini–Hochberg method. E) *proLOX2::GUS* staining following wounding and lanolin treatment, showing that this induction of GUS activity was not blocked by lanolin treatment. ns: not significant with unpaired (Welch’s) t-test. Each dot represents a measurement from one wounded leaf. F) Expression of *COR15A* in *aba3-1* mutant. ns: not significant with unpaired (Welch’s) t-test. Each dot represents a biological replicate.

**Figure s7.**
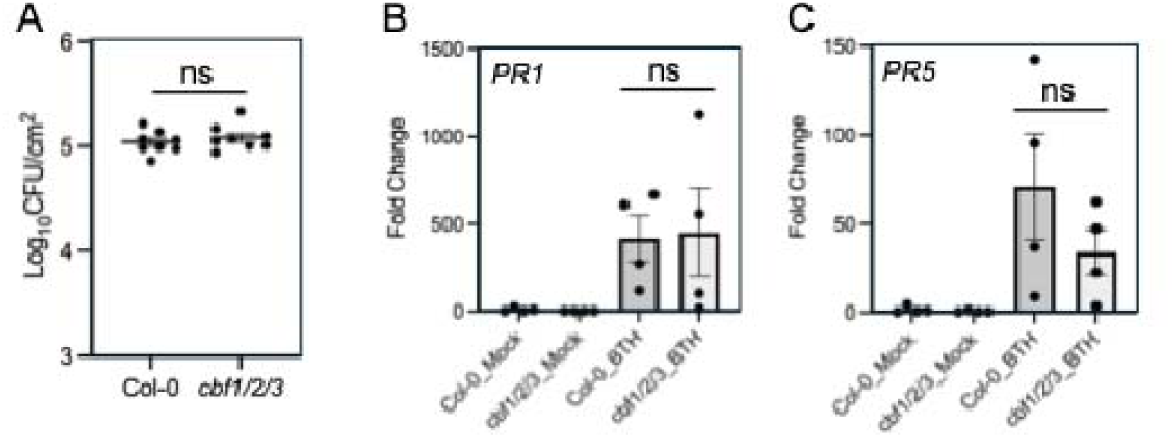
CBFs mutations did not alter resistance to bacterial pathogens and BTH-induced SA activation. A) Bacterial growth in Col-0 and the *cbf1/2/3* mutant at 3 days post inoculation. B–C) Induction of *PR1* and *PR5* in Col-0 and *cbf1/2/3* mutants under mock or BTH treatment. BTH was applied by foliar spray, and no wounding occurred during the treatment. Ns: not significant using unpaired t-test with a two-tailed hypothesis. Each dot represents a biological replicate.

**Supplementary Table 1: List of genes Figure 1A and Figure 6A**.

**Supplementary Table 2: Numeric raw data.**

